# Preprint peer review enhances undergraduate biology students’ disciplinary literacy and sense of belonging in STEM

**DOI:** 10.1101/2022.10.06.511170

**Authors:** Josie L. Otto, Gary S McDowell, Meena M. Balgopal, Rebeccah S Lijek

**Author notes:** **Source of Support:** National Science Foundation Award 2142108: Collaborative Research: Developing Biology, Undergraduates’ Scientific Literacy and Identity Through Peer Review of Scientific Manuscripts.

## Abstract

Undergraduate education on science publishing and peer review is limited compared to the focus on experimental research. Since peer review is integral to the scientific process and central to the identity of a scientist, we envision a paradigm shift that makes teaching peer review integral to undergraduate science education, and hypothesize that this may facilitate the development of students’ scientific literacy and identity formation. To this end, we developed a curriculum for biology undergraduates to learn about the mechanisms of peer review, then write and publish their own peer reviews as a way to authentically join the scientific community of practice. The curriculum was implemented as a semester-long intervention in one class and as a module intervention embedded into a discipline-based class on vaccines. Before and after both interventions, we measured students’ scientific literacy, including peer review ability, using quantitative methods. We also carried out a longitudinal qualitative assessment of students’ perceptions of their scientific literacy and identity using thematic analysis of students’ writing. Here, we present data on the improvement in peer review ability of undergraduates in both classes, and data on the curriculum’s interrelated impact on students’ development of scientific literacy, identity, and belonging in academic and professional spaces. These data suggest that undergraduates can and should be trained in peer review to foster the interrelated development of their scientific literacy, scientific identity, and sense of belonging in science.

## Introduction

Undergraduate science education often focuses on how experiments are carried out and the knowledge generated by the resulting research literature, but misses an opportunity to engage students in the critical validation process that translates one into the other - the peer review of primary scientific literature. Authentic laboratory experiences (e.g. CUREs, independent research) are becoming recognized as important to undergraduate science education as they enculturate students in the science community and increase understanding about the principles of experimental research (1). The value of early research experiences depends on them being authentic and within a community of practice (CoP, see **Table 1** for a definition of terms (1–3)). Moreover, implicit teaching of scientific inquiry through experimentation is insufficient for students to learn how scientists engage in inquiry (4, 5). As a result, scientists’ experiences of science, and what students in classrooms learn about, and conceive of, the process of science, are demonstrably different (6–8). Missing from many undergraduate research experiences are opportunities to learn how scientists *communicate* through scholarly publishing and peer review. We posit that providing undergraduates with explicit instruction in authentic forms of scientific communication - like peer review - will develop their scientific literacy and disciplinary literacy (**Table 1**).

**Table 1:** Definitions of terms used in this manuscript.

| **Term** | **Definition** |
| --- | --- |
| **Community of practice** | A model for studying learning and identity development of individuals who share ways of thinking, communicating, or doing as they develop mastery of knowledge and skills through participation in the community (1). |
| **Scientific literacy** | The ability to know how scientific knowledge is generated and used to make evidence-based claims, and how to make authentic scientific content. |
| **Disciplinary literacy** | What it means to think, read, communicate, and use information like an expert in a particular discipline |
| **Authentic peer review** | The process of writing critiques of scientific research manuscripts to evaluate and improve their scientific integrity and clarity. We use the modifier “authentic” to distinguish this process from when students evaluate *other students’ classwork*, which is not the focus of this study. |
| **Constructivist** | An approach to education based on learning through experience, which acknowledges that learning is an active and socially-constructed process. |
| **Service-learning** | The integration of academic activities with community needs, combining service with reflection in a structured learning environment. |
| **Preprint** | A scientific research manuscript that the authors openly share on a free, online server, usually prior to journal-organized peer review and curation. |
| **Scientific identity** | The composition of self-views as someone who knows about, uses, and contributes to science as part of the scientific community. |

One way in which undergraduates can learn about and engage in authentic scientific conversation with a community of practicing researchers is by participating in the peer review of manuscripts. Peer review is integral to the scientific process, yet authentic scholarship experiences for undergraduates (such as writing and publishing manuscript reviews) are rare. The process of peer review is the backbone of scientific inquiry and a central component of the identity of a scientist (9). It justifies public confidence in scientific results and drives decisions about what research is published and funded. Therefore, education about peer review, and participation in authentic peer review (**Table 1**), ought to occupy a central role in undergraduate science education, in the same vein as education about experimental research and participation in laboratory research (10, 11). Peer review is a form of disciplinary literacy (**Table 1**) and as such peer review is often reserved for those perceived as experts (e.g. faculty) and explicitly excludes students (12, 13). Yet, how can students develop disciplinary literacy - what it means to think, read, communicate, and use information like an expert in a particular discipline - if they are not provided instruction and authentic practice in this essential skill? Therefore, peer review represents part of STEM’s “hidden curriculum” of unstated norms, values, skills, and expectations that are untaught yet required for success (14). Since peer review is integral to the scientific process and central to the identity of a scientist, we envision a paradigm shift that makes *teaching* peer review integral to undergraduate science education (15). Just as early research experiences help students form a scientific identity (16) and develop scientific literacy (17), so too could early scholarship experiences in peer review.

To this end, we developed and assessed a constructivist, service-learning curriculum for undergraduates to learn about the mechanisms of peer review, then write and publish their own peer reviews as a way to join the scientific community of practice. This contrasts with the traditional didactic model of peer review education currently practiced in laboratories. Our recent analysis of early career scientists revealed that formal, evidenced-based instruction in peer review is rare (12) and so there is an unmet need to develop curricula on this topic. The curriculum was designed with the goal of positively contributing to student learning, pedagogical research, and society (see **Figure 1**). It leveraged an innovation in scientific publishing, preprints, which are scientific manuscripts uploaded by the authors to a free, public server often at the same time as submission to a peer reviewed journal (18). Depositing articles as preprints on servers prior to journal submission has long been a normal practice in fields such as physics and mathematics, and has recently grown in popularity in the biological sciences (19, 20). At the same time, experiments in open and pre- and post-publication peer review (21–23) have created preprint review platforms such as Review Commons (24), Early Evidence Base (25), Sciety (26), and PREreview (27). These platforms remove peer review from the exclusive realm of journals to increase participation in the peer review process. In contrast to participating in traditional journal club activities using already finalized and published journal articles, undergraduate students now have the opportunity to engage in authentic peer review experiences, see work in progress, and experience the joy of working to improve the integrity and clarity of scientific manuscripts.

**Figure 1:**
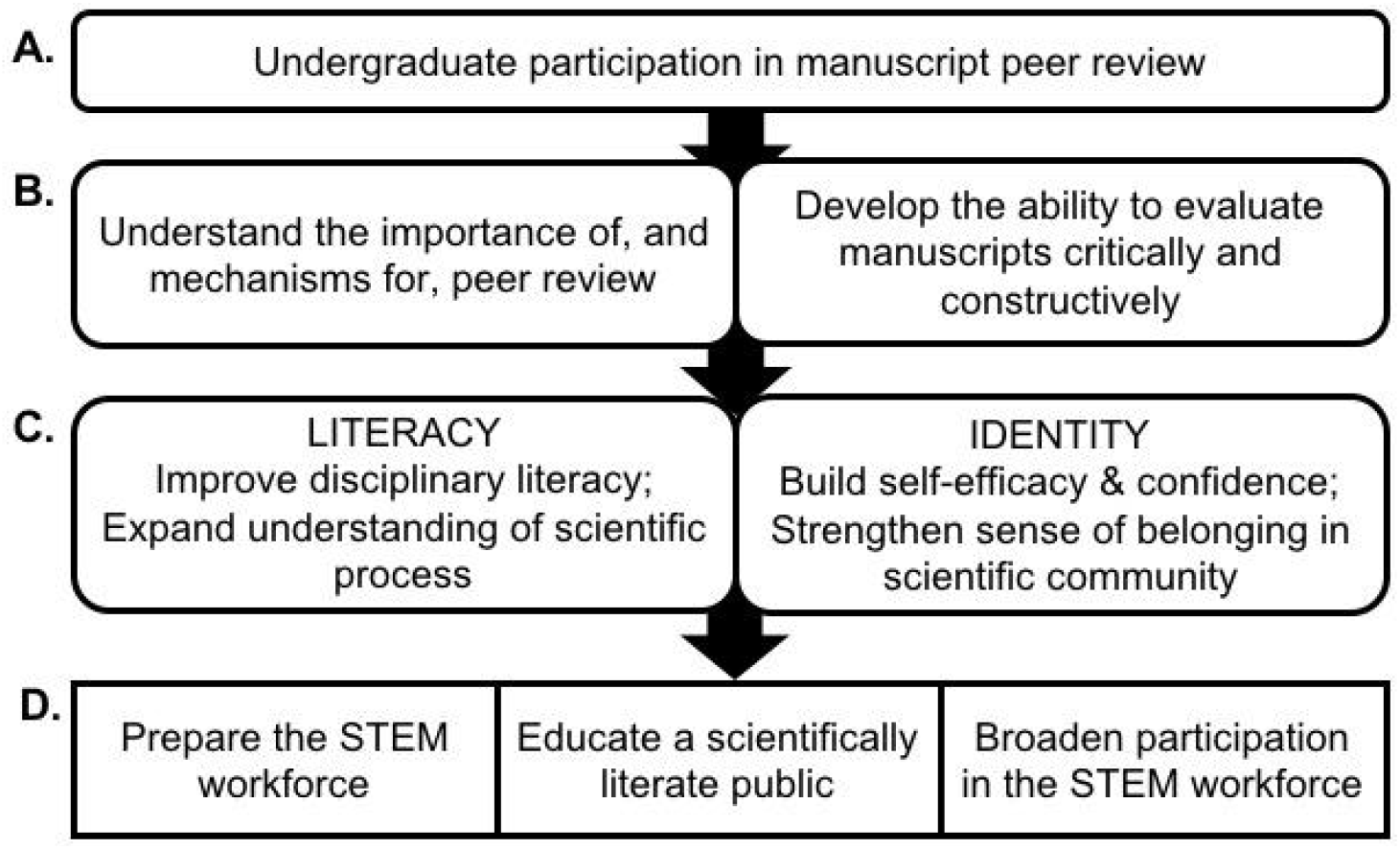
Proposed impact of the curricular intervention (A) on student learning goals (B), research outcomes (C), and benefits to society (D). By explicitly teaching students about peer review and engaging them in it, we hypothesized that this curriculum would develop students’ disciplinary literacy (the ability to think, read, communicate, and use information like an expert in a particular discipline) and scientific identity (the composition of self-views as someone who knows about, uses, and contributes to science as part of the scientific community).

Using preprints instead of published primary literature circumvents the problem that engagement with published literature overlooks the authentic growth processes that occur prior to publication. It gives students a retrospective narrative of scholarship, instead of the more realistic view of science as a constant work-in-progress in which failures and corrections are common. Undergraduates may struggle to reconcile final polished work with their personal experiences with science: failed experiments, negative data, disproven hypotheses. This disconnect could in turn negatively affect their sense of belonging in science and their understanding of the nature of science. Another advantage to using preprints - live, first-draft manuscripts - is that it gives students an opportunity to authentically help real scientists improve their work by sharing their reviews with the authors. Students had the opportunity to publish their peer reviews of preprints on open-access, journal-independent internet platforms as a way to authentically engage with the scientific literature and the scientific community of practice.

Using an apprenticeship model to explore an undefined “space of learning” (16), the curriculum aimed to facilitate students’ self-development and self-expression within a community of practice, both within an academic setting (e.g. by engaging with peers in the classroom) and a professional setting (e.g. by engaging with preprint authors by publishing reviews in professional online forums). Here, we present findings from an exploratory study that used mixed methods to the interrelated impact of our curricular intervention on students’ sense of science identity, literacy, and belonging in academic and professional spaces. We hypothesized that this curriculum, by explicitly teaching students about peer review and authentically engaging them in a community of practice, would improve students’ disciplinary literacy (specifically, peer review ability) and foster a sense of scientific identity and belonging in the scientific community.

## Methods

### Context: Intervention and Participants

In a research liberal arts college for female, transgender, and non-binary students (Mount Holyoke College), we implemented a curriculum on peer review in two different contexts: a) as a full 14-week seminar course (Course 1: Peer Review in Biology) or b) as a single unit of peer review activities embedded within a disciplinary biology course (Course 2: Vaccines). Both courses were offered as upper-level electives and taught by the same instructor (RSL) in the same semester (Spring 2022). In Course 1, with nine students, peer review activities were scaffolded to transition the student from a novice to an expert through four units, based on the clinician training paradigm of “see one, do one, teach one” (28) and the gradual release of responsibility model of literacy education that uses the framing “I do, we do, you do” (29). **Figure 2** provides a conceptual overview of the full curriculum taught in Course 1. In Course 2, with 11 students, only one minimal unit (i.e., “do one,” where students perform peer reviews) was implemented to complement a discipline-specific course on vaccine biology. In both classes, the instructor and/or students selected biology preprints of interest to review, critically analyzed the preprints in writing and in discussion using guiding questions provided by the instructor, and then wrote peer review reports as a professional peer reviewer would do (see “Context: Peer Review Activities” below for further detail). Students submitted weekly reflection journals in response to prompts about their perceptions of their performance, sense of self-efficacy, and understanding of disciplinary literacy in the context of peer review. A majority of students indicated their intention to publish their reviews publicly online to document their expertise and participate in the professional CoP. Students also interacted with preprint authors and other experts in peer review through email and video interviews. All students provided informed consent to participate in the study, which was verified by the Mount Holyoke Institutional Review Board as Exempt according to 45CFR46.101(b)(1, 2): (1) Educational Research, (2) Tests, Surveys, Interviews on 10/12/2021.

**Figure 2:**
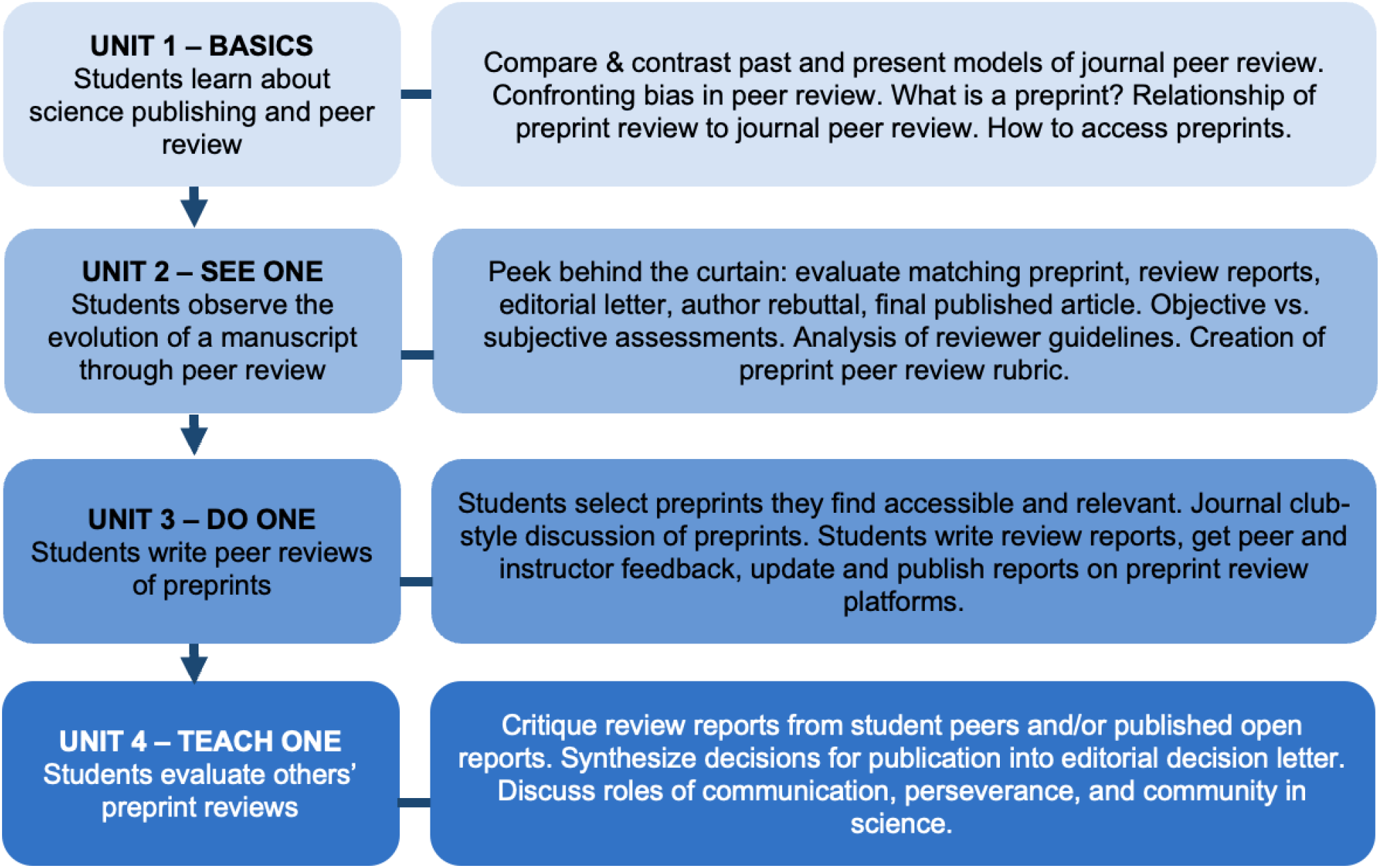
Peer review curriculum. In this constructivist service-learning curriculum, peer review activities are scaffolded to transition the student from novice to expert through 4 units, loosely based on the clinician training paradigm of “see one, do one, teach one” (28) and the gradual release of responsibility model of literacy education that uses the framing “I do, we do, you do” (29). Educators could choose to use unit(s) alone or together depending on course needs and students’ previous experience. Throughout the curriculum, students review preprints freely available on servers and have the opportunity to publish their reviews to document their scholarship and serve the scientific community.

### Context: Curriculum and Peer Review Activities

Course 1: Peer Review in Biology students engaged in four peer review events throughout the full curriculum (**Figure 2**):

1. *Review 1: Individual review on a manuscript selected by the instructor*. The initial, baseline event was assigned as individual homework after the first class meeting, after an initial discussion of the concept of peer review but before students had carried out any in-depth training in the course. All students reviewed the same manuscript, selected by the instructor for its accessibility to a general biology audience. The manuscript was written by pre-college students, submitted to the *Journal of Emerging Investigators* (30), and ultimately published as (31).
2. *Review 2: Individual review on a preprint selected by student groups*. The next event took place in Unit 3, after ~6 weeks of explicit teaching about peer review and after students co-created a rubric to evaluate preprint peer reviews (32, 33). Students were grouped by interest in biological topics (e.g. cancer, COVID19) and then each group selected one preprint to review, which was approved by the instructor as being accessible. Each group member individually carried out peer review of the same preprint as homework. Students were *not* provided with detailed written instructions for how to write a peer review; instead they were asked to create their review based on their learning from Units 1 & 2. Guiding questions for peer review were provided as an optional resource but answers were not required to be provided in the assignment.
3. *Group Review 2: Group review on a preprint selected by student groups.* Students shared their individual Review 2 with their group members, then spent time in class discussing their individual reviews and the instructors’ feedback on them. Then they synthesized their individual reviews into one group review (the third review event). After completing the group review, students re-read their individual reviews and self-graded using the rubric they co-created (32, 33).
4. *Review 3: Individual review on a preprint selected by individual students*. The fourth review event was assigned as an individual, final assessment after the completion of the curriculum. Without any input from the instructor, each student selected a preprint of interest and wrote an individual review.

Course 2: Vaccines students engaged in a single module of the curriculum (Unit 3, **Figure 2**) and three peer review events:

1. *Review 1: Individual review on a manuscript selected by the instructor*. The initial, baseline event was assigned as individual homework after a 30-minute discussion of the concept of peer review and after ~6 weeks of disciplinary lessons on vaccinology. All students reviewed the same preprint, selected by the instructor for its accessibility to a general vaccinology audience (34). Because this course did not involve explicit lessons on peer review in class, students were instead provided with detailed written instructions for how to perform a peer review, including guiding questions which were required to be answered as part of the assignment.
2. *Review 2: Individual review on a preprint selected by student groups*. Students were grouped by interest in vaccine topics and then each group selected one preprint to review, which was approved by the instructor as being accessible. Each group member individually carried out peer review of the same preprint as homework, using the same detailed instructions and guiding questions as Review 1.
3. *Group Review 2: Group review on a preprint selected by student groups*. Students shared their individual Review 2 with their group members, then spent time in class discussing their individual reviews and the instructors’ feedback on them. Then they synthesized their individual reviews into one group review (the third review event).

### Assessment of disciplinary literacy - peer review quality

All peer reviews written by students were de-identified by the instructor (RSL) and provided to the independent researchers (GSM) to assess within-student development of peer review ability longitudinally. The researcher generated four metrics of peer review ability using <u>three</u> unique tools:

1. *The Review Quality Instrument* (RQI, (35) which consists of eight Likert scale questions (ranging 1-5). One question asks the evaluator’s overall opinion of the review, and this question is reported here as *“RQI”* (range = 1-5). The other seven questions ask about components of the review, and here these have been combined and reported as “*RQI Total”* (range = 7-35).
2. *The PREreview Review Assessment Rubric (32)* (32)
3. A rubric for evaluation of preprint reviews was generated by the students in Course 1 (32, 33). This consists of a series of non-linear scores of 0-4 being awarded to different sections of the review, which are then converted into a percentage, reported here as “*MHC*”.

Additionally, scientific literacy more broadly defined was assessed using Gormally’s TOSLS Survey (36) administered before and after each of the two interventions. TOSLS results are presented in Appendix 1.

### Thematic Analysis

Students completed weekly reflection journals, which were de-identified by the instructor (RSL), provided to the independent qualitative researchers (JLO, MB), and assigned pseudonyms to retain anonymity. De-identified reflection journal entries were uploaded to MaxQDA (ver. 22.2.1; 2022) and coded by thematic analysis (37) (**Appendix 2**). Thematic analysis used students’ science literacy, science identity, and belonging within the scientific community to inform initial latent codes. Initial themes corresponding to science literacy included: understanding science content, using science skills to help others, talking about science with others, and practicing science now and in the future. These codes were then collapsed into the following overarching themes: *knowledge* (understanding and talking about science), *practice* (applying/performing science skills and knowledge), *and value of practice* (understanding the use and need of science). We also identified personal and environmental variables corresponding to a student’s identity that were then organized as *professional identity* (pursuing a science career or internship and interacting with science professionals) and *personal identity* (systemic/structural barriers and access to resources). Finally, we divided codes relating to belonging into *presence, absence, or facultative* (i.e., a sense of belonging in some contexts but not others). To identify differences between each course context, we compared codes across courses. To establish trustworthiness, two team members separately coded 20% of the students reflections and ensured a minimum 90% inter-rater reliability. We also engaged in expert debriefing to discuss the alignment between our themes and our theoretical framing.

### Inclusion and exclusion criteria

Only data from students who completed all assignments were included in the analysis. This represents all 9 students in Course 1, and 10 out of 11 students in Course 2. In Course 2, one student did not complete reflection journal entries and so that student’s data was excluded from the thematic analysis. Another student did not complete the peer review exercises on time nor the post-survey for TOSLS, and so this student’s data was excluded from those analyses.

## Results

### Assessment of Disciplinary Literacy by Evaluating Peer Reviews

The impact of the curricular interventions on students’ peer review ability - a form of disciplinary literacy - was measured quantitatively by an independent researcher using three unique instruments. Students’ peer reviews from both courses were de-identified by the instructor (RSL), and their quality was assessed by the independent researcher (GSM) using the Review Quality Instrument (RQI, (35)), the PREreview Review Assessment Rubric (32) and a rubric that was created by the students themselves (in Course 1, “MHC,” (32, 33)). **Table 2** and **Figure 3A** summarize the results of the four peer review events in the full peer review curriculum (Course 1). **Table 3** and **Figure 3B** summarize the results of the three peer review events in the peer review module that was embedded into a disciplinary course (Course 2, see **Methods: Context - Curriculum and Peer Review Activities** for a description of each peer review event). Students in the full curriculum (Course 1) showed increases in peer review ability after each peer review event using all 3 evaluation tools (**Table 2, Figure 3A**). Students in Course 2 showed an increase in peer review quality throughout the duration of the course using all 3 evaluation tools, mostly reflected in the transition from individual reviews to group reviews (**Table 3, Figure 3B**). Overall and depending on the measurement tool used, the full curriculum resulted in a 25-42% increase in students’ peer review ability (**Table 2**, bottom row), which was comparable to the 14-40% improvements seen as a result of the embedded module (**Table 3**, bottom row). Measurement of improvement was lowest on the PREreview assessment because many reviews were close to the ceiling on this tool. At the final peer review event, which occurred after completion of the interventions, students’ reviews in both classes earned 80-97% of the total maximum score possible for each assessment tool, remarkable because the RQI and PREreview tools were designed to assess the quality of reviews written by *experts*, not novices or other learners. Many students elected to publish their reviews (e.g. (38–42)), implying that they were proud of the final products (see Assessment of Students’ Perceptions, below). These data suggest that while the baseline quality of peer reviews written by an untrained undergraduate (novice) is mediocre as might be expected, these disciplinary literacy skills can be developed through an intentional curriculum that offers explicit instruction, iterative practice, and opportunities to authentically participate in a community of practice.

**Table 2:** Changes in peer review quality as a result of the full curriculum (Course 1). Students carried out 3 individual reviews as described in the Methods, with Review 2 being used to generate the Group Review. Changes in scores between different reviews are reported as Δ(Review#);(Review#), and are normalized to give a range of change from −1 (maximum decrease) to +1 (maximum improvement), with 0 signifying no change in peer review quality. Review quality was assessed by an independent evaluator using four metrics: RQI: the single question in the Review Quality Index where the evaluator gives an overall assessment of the review; RQI total: the combined score all questions in the RQI; PREreview Reviewer’s Assessment Rubric score; and MHC: the grading rubric created by students in Course 1.

| Tool |  | RQI | RQI Total | PREreview | MHC |
| --- | --- | --- | --- | --- | --- |
| Range |  | [1-5] | [7-35] | [1-5] | [0-100%] |
| Average score | **Review 1** | 2.33 | 19.4 | 3.52 | 50.8% |
|  | **Review 2** | 3.22 | 21.1 | 4.18 | 73.2% |
|  | Group Review | 3.56 | 23.7 | 4.33 | 81.0% |
|  | **Review 3** | 4.33 | 28.1 | 4.84 | 92.9% |
| Change per event | $\Delta(1\blacktriangleright 2)$ | 0.18 | 0.05 | 0.13 | 0.22 |
| | $\Delta(2\blacktriangleright 3)$ | 0.22 | 0.2 | 0.13 | 0.2 |
| | $\Delta(1\blacktriangleright 3)$ | **0.4** | **0.25** | **0.26** | **0.42** |

**Figure 3:**
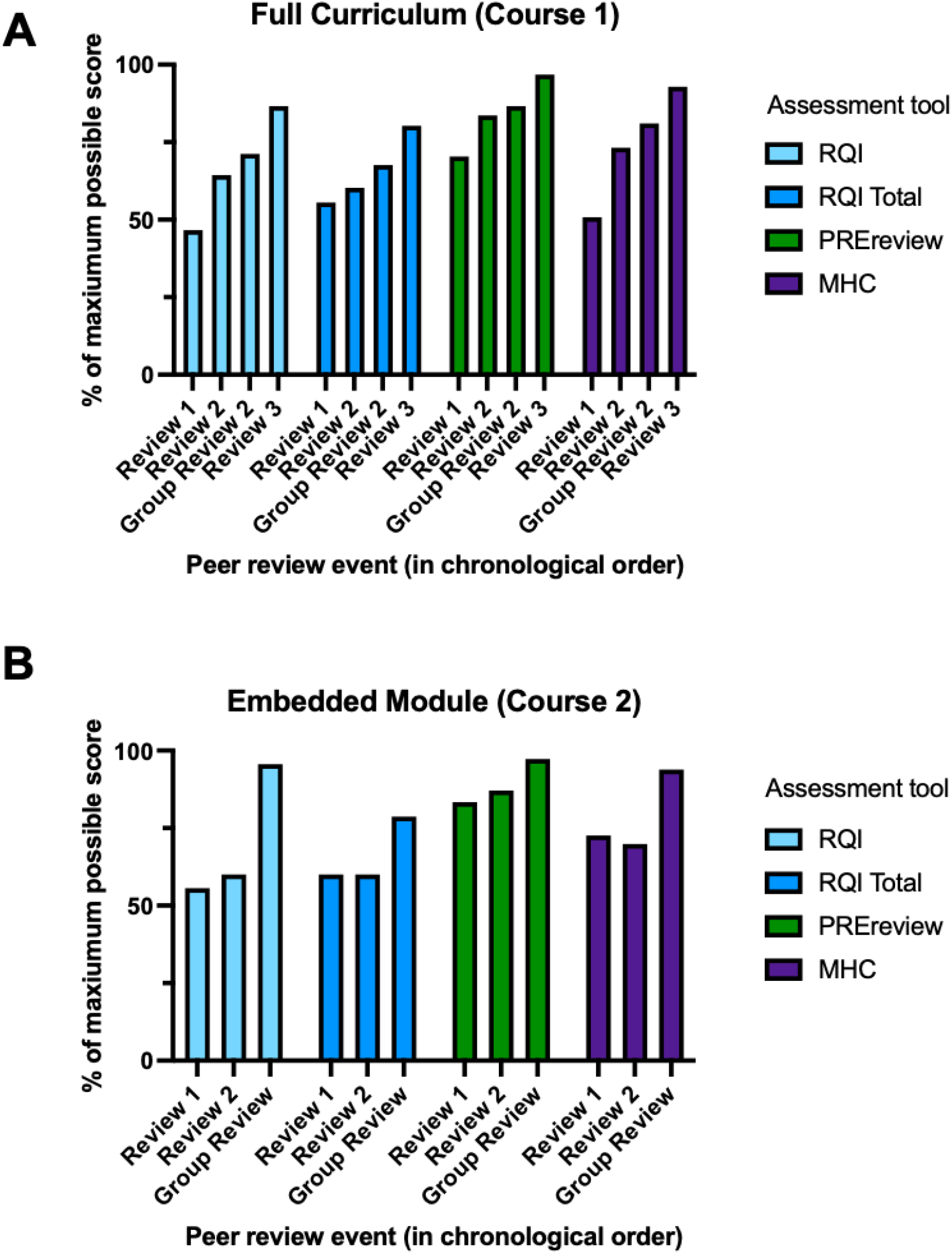
Improvements in peer review quality as a result of the full curriculum (A) and embedded module (B). Students’ de-identified peer reviews were assessed by an independent researcher using four metrics: RQI: the single question in the Review Quality Index where the researcher gives an overall assessment of the review; RQI total: the combined score all questions in the RQI; PREreview Reviewer’s Assessment Rubric score; and MHC: the grading rubric created by students in Course 1. Data are presented as a percent of the maximum score possible on each of the tools to compare improvement across tools. Review events are presented on the x-axis in chronological order in the curriculum.

**Table 3:**
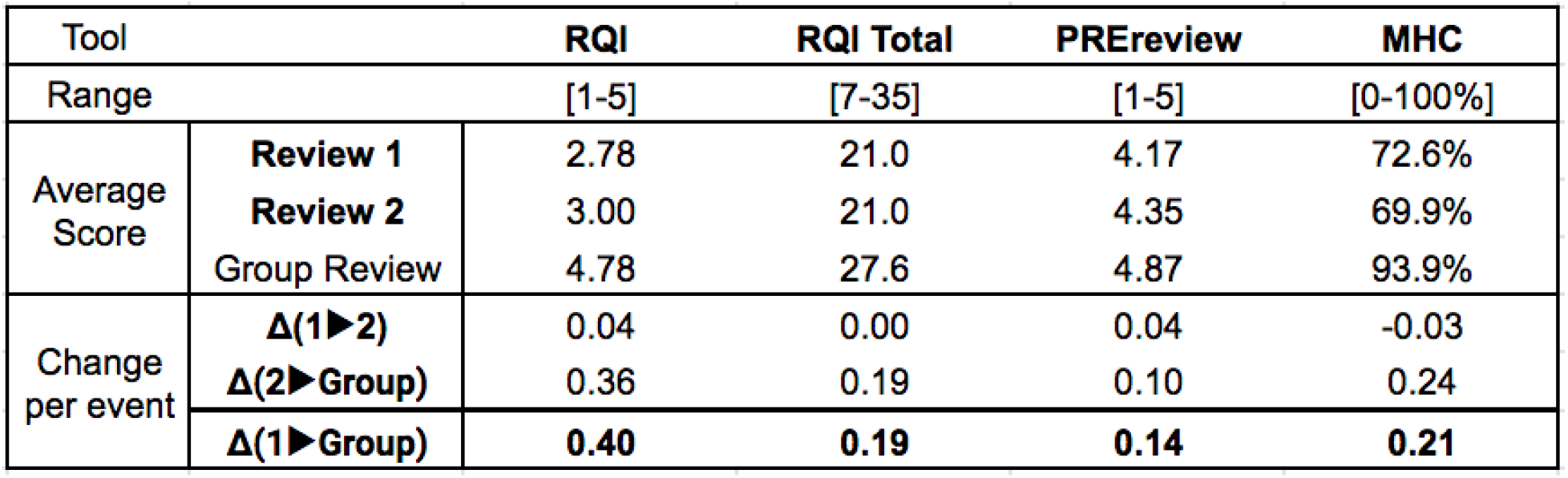
Changes in peer review quality as a result of the embedded module (Course 2). Students carried out 2 individual reviews as described in the Methods, with Review 2 being used to generate the Group Review. Changes in scores between different reviews are reported as Δ(Review#);(Review#), and are normalized to give a range of change from −1 (maximum decrease) to +1 (maximum improvement), with 0 signifying no change in peer review quality. Review quality was assessed by an independent evaluator using four metrics: RQI: the single question in the Review Quality Index where the evaluator gives an overall assessment of the review; RQI total: the combined score all questions in the RQI; PREreview Reviewer’s Assessment Rubric score; and MHC: the grading rubric created by students in Course 1.

### Assessment of Students’ Perceptions of Science Literacy, Identity, and Belonging by Thematic Analysis

Students’ perceptions of their own development were captured in both courses through weekly reflection writing in response to specific prompts. Reflections were de-identified by the instructor and provided to the independent evaluators (JLO, MB) for thematic analysis. Results from this analysis demonstrated that students developed an affiliation with the science CoP over the course of the interventions. While many students were science majors (hence, early career scientists), a few used the course as an opportunity to engage in and learn about science as non-practitioners. In this sense, students already identified as “novices” within the CoP or transitioned from that of an “outsider.” Through participating in the peer review curriculum, students reflected on their literacy, identity, and belonging within the CoP. **Table 4** summarizes whether each student showed evidence of these themes by the conclusion of the interventions, and the codebook used for thematic analysis, along with exemplary quotes from students, is in **Appendix 2**.

**Table 4.** Componential analysis of the overarching themes of literacy, identity, and belonging achieved by students upon completion of the intervention, either after the full curriculum (Course 1) or the embedded module (Course 2).

|  | Students | Science Literacy<sup>1</sup> |  |  | Science Identity |  | Science Belonging<sup>1</sup> |  |
| --- | --- | --- | --- | --- | --- | --- | --- | --- |
|  |  | Knowledge | Practice | Value | Personal<sup>2</sup> | Professional<sup>1</sup> | Academic | Professional |
| Course 1 | Kasper | NO | ✓ | ✓ | -,+ | ✓ | ✓ | NO |
|  | Kori | ✓ | ✓ | ✓ | -,+ | ✓ | ✓ | ✓ |
|  | Gianna | NO | ✓ | ✓ | - | ✓ | ✓ | NO |
|  | Niki | ✓ | ✓ | ✓ |  | ✓ | ✓ | ✓ |
|  | Dani | ✓ | ✓ | ✓ | + | ✓ | ✓ | ✓ |
|  | Aaliyah | ✓ | ✓ | ✓ |  | ✓ | ✓ | ✓ |
|  | Kiara | ✓ | ✓ | ✓ | - | ✓ | ✓ | ✓ |
|  | Zara | ✓ | ✓ | ✓ | - | ✓ | ✓ | ✓ |
|  | Madi | ✓ | ✓ | ✓ | - | ✓ | ✓ | ✓ |
| Course 2 | Sam | ✓ | ✓ | ✓ |  | ✓ | ✓ | ✓ |
|  | Alex | NO | ✓ |  |  | ✓ | ✓ | NO |
|  | Amara | ✓ | ✓ | ✓ | - | ✓ | ✓ | ✓ |
|  | Isabella | ✓ | ✓ |  |  | ✓ | ✓ | NO |
|  | Sofie | ✓ | ✓ | ✓ |  | ✓ | ✓ | ✓ |
|  | Kris | ✓ | ✓ | ✓ | + | ✓ | ✓ | ✓ |
|  | Riley | ✓ | ✓ |  |  | ✓ | ✓ | ✓ |
|  | Jocelyn | ✓ | ✓ | ✓ |  | ✓ | ✓ | ✓ |
|  | Aneta | ✓ | ✓ | ✓ |  | ✓ | ✓ | ✓ |
|  | Lotte | ✓ | ✓ |  |  | ✓ | ✓ | ✓ |

<sup>1</sup> Checkmarks (✓) indicate which students were coded as being scientifically literate, possessing a professional science identity, or having a sense of belonging within the science community.
(NO) indicates students who were coded as not possessing scientific literacy, identity, or belonging.
Cells are left empty when student responses don't address aspects of literacy, identity, or belonging.
<sup>2</sup> (-) indicates aspects of a student's personal identity that the student perceives as a disadvantage.
(+) indicates aspects of a student's personal identity that the student perceives as an advantage.

#### Literacy

In both courses, the peer review curriculum improved students’ perceptions of their own scientific literacy. Scientific literacy included the students’ understanding of how scientific knowledge is generated, their engagement in science practices, and how they perceived the value of these practices. For example, students coded as scientifically literate were able to describe the peer review process and its value, as well as feel confident about performing a peer review. This is seen in Sam’s journal entry when she commented on her confidence in performing peer review after the intervention: “*After thoroughly reviewing the standard requirements for a publication, as well as the scientific theory supporting the article, I felt surprisingly well-equipped to offer constructive feedback on the assigned pre-print*.”

A nuanced development and then strengthening of scientific literacy was seen in the full curriculum (Course 1) over each of the four units (**Figure 2**). In Units 1 and 2, students primarily reflected on their new understanding of what peer review is, who engages in peer review, and how to do one. In their early journal entries, students shared many of the misconceptions that they held about peer reviews, including that reviews were done after publication and that authors were responsible for recruiting reviewers themselves. This is seen in Kiara’s reflections when she states: “*I had this idea that there was not an editor involved, but rather that the author of the study themselves was responsible for reaching out and facilitating the process*.” Students went on to refute these misunderstandings, even explaining why they were wrong. Within this unit, students also shared their perceived value of the peer review process. Most students suggested “superficial” benefits such as the ability to edit articles prior to publishing them, but did not yet demonstrate a sophisticated understanding of the value of peer review plays for science (43). In Unit 3, during which students completed Review 2, the first review after instruction, (see **Methods - Context: Curriculum and Peer Review Activities**), all students discussed feeling confident, comfortable, or prepared for the assignment. Students explained that the course activities and materials helped to clarify what was expected of their review. In Unit 4, when students evaluated others’ reviews, there were obvious improvements in students’ depth of knowledge of the value of peer review. Several students were able to elaborate on the importance of reviewers in producing high quality publications. Kiara shared: “*This feature is incredibly valuable to the process as it allows for the most amount of feedback for the author … as the more feedback that is provided for the review, the better the review can potentially be.”* Kiara’s comments demonstrated an understanding that the review process contributes to how scientific knowledge is disseminated. Students in Course 2 were also seen to strengthen their literacy through critique of discipline-specific papers (e.g. on vaccines) and offering constructive feedback on peer presentations that occurred outside the embedded peer review module. Lastly, students in both courses mentioned that through reading and critiquing reviews made by their peers, they were able to sharpen their own skills in preparation for their finals reviews. Many students discussed their plans for their final review, each noting unique areas of improvement. Kiara was inspired by her classmates’ use of a peer review rubric to keep track of necessary comments, stating that she would remember to use it in the future. Over the course of the semester, students first developed their knowledge, value, and practice of peer review and then further strengthened these points through repeated writing and critiquing of reviews.

#### Identity

In both courses, students’ perception of their science identity was influenced by their own personal and professional identities. Students who demonstrated a strong science identity talked about science with others in and out of class, described the ways in which their newly learned skills could be applied to their academic and post-graduate careers, and expressed their validation in the science field because of others who looked like them. Madi wrote: “*the critical thinking skills I gained from this peer-review course have really started kicking in. I also feel confident that if my experiment works, I will be able to write about it for a manuscript.*” Earlier in her reflection, Madi explained how learning to analyze and critique research data helped her to overcome a roadblock in her research lab. “*I am also considering writing a thesis for my senior year because of the confidence I have gained in research, reading, writing, and reviewing.*” Madi demonstrated that through reflection of her own experiences, she planned to engage in future disciplinary literacy activities that will likely further solidify her science identity.

#### Belonging in the academic CoP

Science literacy and identity within the CoP facilitated a student’s progression from “novice” to “expert” when they felt they belonged (**Figure 4**). We identified two “spaces” in which students felt they belonged. The academic space consisted of in-person and virtual classrooms, where students interact with one another. The professional space is where students interact with science professionals (e.g. researchers, journal editors, expert reviewers). Engaging with classmates through discussion and co-writing peer reviews reinforced students’ sense of belonging within the academic space. A few students felt they belonged in some contexts, but not in others. For example, Isabella explained, “*I think discussing the paper in class with people who are on the same level as I really helped*.” Here, Isabella explains that the context of a classroom feels safe, but she goes on to explain that in other contexts, where she perceived that there were some members who were different from her, she did not feel as confident. “*In the past, I have done journal clubs with graduate students, which have been much more difficult for me to feel confident in sharing my thoughts. Today’s class really helped me gain more confidence about my ability to review primary research independently*.” Hence, some students, like Isabella, expressed confidence in academic spaces but not in professional ones prior to the intervention, and implied that the confidence gained through the intervention could improve her feelings of belonging in professional spaces in the future.

**Figure 4:**
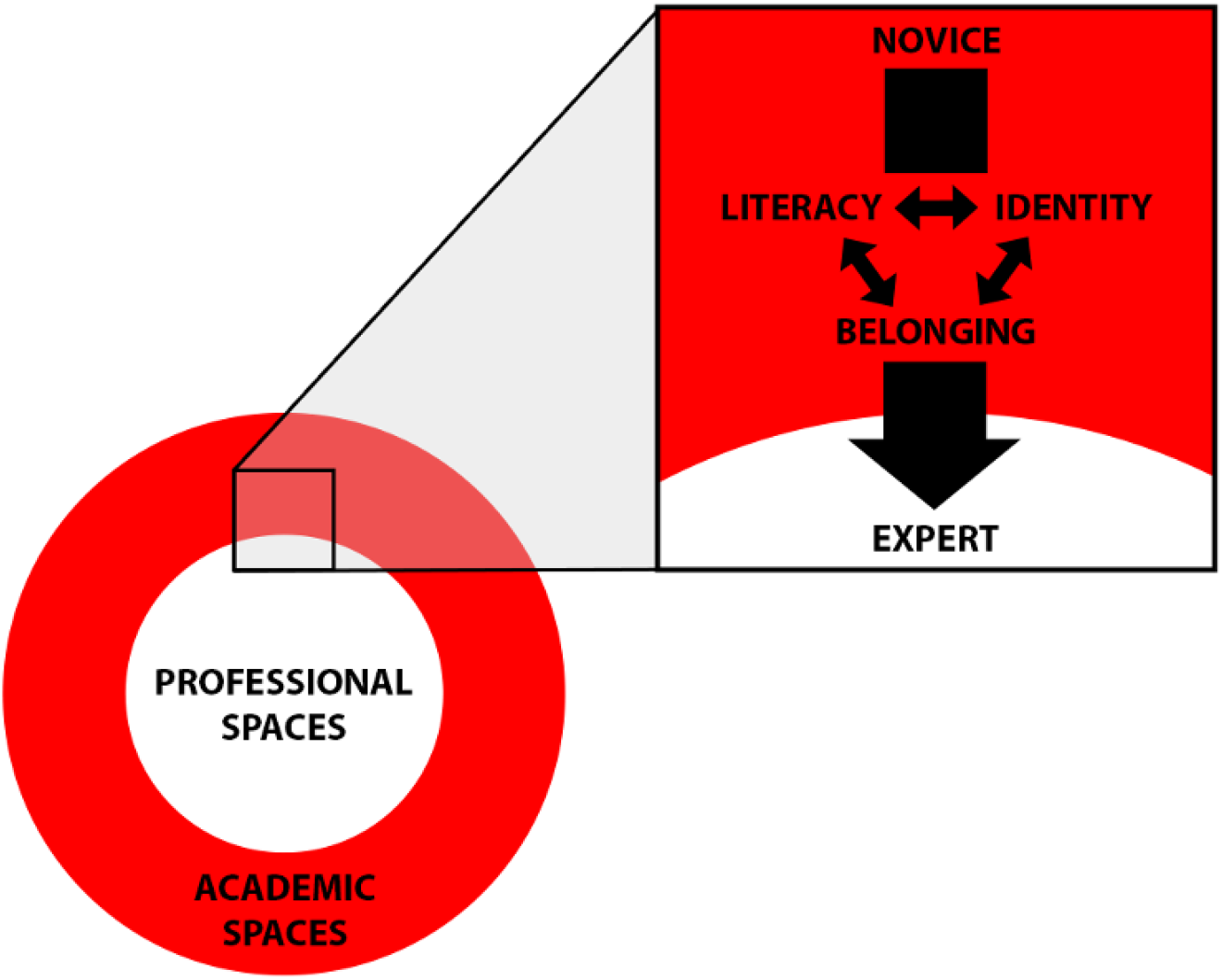
Community of practice (CoP) conceptual model (adapted from (1)). Students’ progression from “novice” to “expert” is influenced by their sense of belonging within both academic and professional spaces. We propose that the “expert” and “professional” spaces align, due to the ability to participate in authentic peer review processes and interact with professional scientists. Belonging is seen to bridge literacy (knowing and applying science) and identity (feeling like a scientist).

#### Barriers to belonging

Students in both courses most often identified illiteracy (i.e., inadequate knowledge and practices) as a barrier to belonging. This is evident in Alex’s journal entry when they mentioned feeling like a “wallflower,” or excluded, because of their lack of science experience: *“I still don’t really feel I am part of the scientific community. It is a huge field and looking at a few papers makes me feel like I am more of a wallflower than anything else… with my very limited experience I don’t think it would be right to say that I am part of the community. I am still going through the initiation rites*.” Although Alex described feeling like they are on the periphery of the CoP, they describe their experience as part of an “initiation” period, which implies that they may see themselves as belonging sometime in the future.

#### Belonging in the professional CoP

Talking with authors and interviewing professional scientists during the interventions allowed for students in both courses to feel they belonged in the professional space (i.e., transition from novice to expert, see **Figure 4**). Madi described how experts helped to foster her belonging within a group of scientists when she wrote: “*…firstly I was able to make a conversation with a professional and deliver my question in a way that did make sense to them. The points they were making made me feel familiar with other jargon and issues surrounding the process of peer review. … I did not feel inadequate or less knowledgeable during these conversations*.” Madi found the experience of interacting with others valuable in shaping her sense of being a member of a CoP. Likewise, Kiara wrote in a journal entry about how she perceived others to perceive her: “*She [professional scientist] saw me as a student and as a scientist.… I so often do not feel seen or understood in the stem field so having this moment to talk to her allowed for me to feel valid in our field*.” In other words, Kiara’s sense of belonging was strengthened through positive reinforcement by those who she saw as experts in the professional science CoP. In their responses, Madi and Kiara pointed to their science literacy and identity being validated through engaging with experts within the CoP.

## Discussion

This study provides evidence that one aspect of disciplinary literacy - peer review of manuscripts - can be developed and strengthened by a curriculum that offers explicit instruction, iterative practice, and opportunities to authentically participate in a community of practice. Improvements in disciplinary literacy and in students’ perceptions of literacy, scientific identity and belonging in STEM were observed as a result of both a standalone peer review course (full, 14-week curriculum) and a short peer review module embedded into a disciplinary course. This suggests that instructors who are not able to offer a full course on peer review might still consider incorporating a module into pre-existing disciplinary courses as a way to intentionally develop students’ disciplinary literacy. Explicit instruction can come in the form of class lessons (as was done in the full curriculum) or written, granular guiding questions (as was done in the embedded module). Interative practice is critical for success (**Figure 3**). The group synthesis (after independent reviews) was particularly beneficial for improving peer review quality, reinforcing why this approach is used in journal peer review when editors synthesize reviews from independent reviewers. Group work also provides students an opportunity to participate in an academic community of practice. Using these pedagogies to foster students’ disciplinary literacy also has interrelated effects on improving students’ sense of belonging in STEM and identity as a scientist. As students transition from novices to experts in their disciplinary literacy skills, it causes them to perceive themselves as valuable members of the academic *and professional* communities of practice, which may explain why these self-beliefs are predictors of persistence in STEM careers, especially for minoritized students (44, 45).

A limitation of our study is that peer reviews are a subjective assessment (of a manuscript), and so evaluations of a peer review are in turn a subjective assessment of a subjective assessment, making it challenging to evaluate peer reviews in a standardized way. This is further complicated where different manuscripts are being evaluated, as the quality of the peer review may well depend on how much (or little) there is to critique in a manuscript. We endeavored to make up for this limitation by using three unique evaluation tools, which are some of the few that are publicly available and allow quantification of some criteria, though there is variability between the tools. None of the tools evaluate the sophistication of the critiques (surface-level comments vs. deep scientific analysis), which we find to be a major limitation yet also difficult to quantify. For example, with the particular preprint chosen for the first review event in Course 2, students had a tendency to request the entire experiment be repeated because of small sample size, but did not state or justify what a suitable expectation for the size would be. Therefore, one future direction of this work is to develop clearer standards for evaluation of peer reviews (and by extension, a rubric for peer review learning outcomes).

All three peer review quality assessments include items to evaluate the “tone” of the review, defining a quality review as having a constructive, respectful tone that balances strengths with weaknesses. This criterion about “tone” speaks to the trend toward overly critical or harsh reviews from professional scientists (12). Interestingly, students’ reviews were never rude or insulting, tended to be written in clear and logical prose, and tended to point out strengths of the works with high frequency, and so consistently scored high marks for these criteria on all three assessment tools. Students’ early career stage and/or gender may have contributed to the civility of their reviews, as both of these identities are socialized to use respectful tone, especially when delivering criticisms. Our sample of students at Mount Holyoke, an institution serving women and minoritized genders, may be biased in this regard. In ongoing work, we are investigating these demographic correlations and curricular outcomes using larger sample sizes in other institutional contexts (e.g. a large land-grant university, a two year college). Regardless, these data suggest that if a balanced approach to reviewing is reinforced at early career stages then it might be better retained in the reviews of professional scientists.

Our data suggest that belonging in a CoP is dependent on the interplay among disciplinary practices, disciplinary discourses, and disciplinary identity (**Figure 4**). Belonging is itself composed of two parts: a sense of belonging in the student’s *academic setting* (i.e. the classroom), in combination with a sense of belonging to the *community of scientists*. The sense of classroom belonging appears to be important in creating a safe space for students to then both develop their literacy and explore their identity as scientists and place in the wider scientific community. Interaction with others - peers, scientists and professionals - then supports the development of the sense of belonging.

Students have a clear understanding of what they are required to do to succeed *academically*, but what *professional* success looks like - and how they can achieve it - may be less clear (46). Addressing in an academic context how to move from novice to expert in an authentic professional practice may be an important foundation in building professional scientific identity. This may have implications for retention in academia: moving instruction on how to be a practicing scientist away from later career stages and into earlier academic settings may allow earlier establishment of belonging in the scientific CoP. Other work has shown that a strong science identity predicts higher grades, with manipulation of belonging in college being able to affect the relationship between science identity and academic performance (45).

Our proposition is supported by prior research on the CREATE method for teaching science, which uses published articles (not preprints) to educate students (47). CREATE students made comments about the importance of personal connections with scientists, which in the original CREATE work, was due to the opportunity for students either to interact with the authors of the published papers that they read, or view materials and footage of a prior group’s interaction (47). These students also both developed literacy about the practices of a CoP, and expressed an increased interest in becoming scientists, identifying with the CoP (47). Similarly, our intervention used primary literature and introduced students to different scientists and practitioners. As in CREATE, we see that when students are given the opportunity to engage with professionals, students’ sense of belonging in the CoP is enhanced, moving them further toward the expert identity. Students also related their science identity to their personal and professional identities, as they described how they communicated about science with friends, family members, classmates, and other scientists. Given the importance of belonging in shaping students’ science identity, and possibly also persistence in STEM (48), one implication of this work is an appreciation of the need for undergraduate biology instructors to understand the nuances between different levels of belonging (46).

Belonging in academic, science, and professional CoPs can all contribute to development of literacy and identity, which then can contribute to a sense of belonging (**Figure 4**). We suggest that undergraduate instruction, especially on literacy development, could potentially be more effective if greater attention is focused on professional belonging. By attending to only one context, instructors may be missing an opportunity for students to identify their own professional goals and whether they believe they can achieve them (44, 45). We posit that undergraduate instructors should design curricula that allow students to reflect on their identities and sense of belonging in both the classroom as well as in the broader science community of practice.

## Acknowledgements

We are grateful to the National Science Foundation for funding this work through an Improving Undergraduate Science Education (IUSE) award from the Division of Undergraduate Education: Award #2142108: Collaborative Research: Developing Biology Undergraduates’ Scientific Literacy and Identity Through Peer Review of Scientific Manuscripts. Portions of that text are adapted here. Our deepest gratitude goes to all of the students who participated in this curriculum and to the colleagues who contributed to it, including Jessica Polka, Daniela Saderi, Sarah Fankhauser, Maisha Miles, and Andrea Henle. Thanks also to the preprint authors for posting their manuscripts, many of whom wrote supportive emails in response to students’ reviews of their work.

