## Supplementary Materials for "Preprint peer review enhances undergraduate biology students’ disciplinary literacy and sense of belonging in STEM"

**Supplemental Materials**

### Appendix 1: Measurement of scientific literacy by TOSLS

We also measured students' broader science literacy using Gormally's Test of Science Literacy Survey (TOSLS, (36)) , which was administered during the first week (pre-survey) and last week (post-survey) of Course 1 and 2. Seventeen questions from the survey were administered in a similar fashion to Cartwright (49). Students in Course 1 scored averages of 76% (13/17, n = 9) in the pre-survey and 69% (12/17, n = 9) in the post-survey. Students in Course 2 scored averages of 72% (12/17, n = 10) in the pre-survey and 81% (14/17, n = 10) in the post-survey. No significant changes in science literacy was apparent in the TOSLS survey data which is likely due to a combination of factors and is not dissimilar to the findings of Cartwright et al. (49), who's method we used. Firstly, the percentage of correct responses closely matches that expected for the kind of population that is at Mount Holyoke College, a private liberal arts college, as demonstrated by Gormally et al. (36). The percentage begins high and so leaves little room for demonstration of improvement. Secondly, the effects of a 14-week class on broad science literacy may well be expected to be minimal, and TOSLS is perhaps more appropriate for longer, even multi-year interventions. However we are continuing testing of the curriculum at other sites, and it may be that TOSLS allows us to see if there are literacy changes in other populations, and allow for some cross-site comparison in future work. It may still be that in populations with a lower initial score in science literacy, engagement with literature through peer review could have the potential to improve competency with scientific content; if it does not, it may be that attention to this aspect needs to be paid when refining the final curriculum at the end of our multi-site evaluation.

**Appendix 2: Thematic Analysis Codebook**

| Theme | Code | Subcode | Definition | Example |
|---|---|---|---|---|
| Literacy | Knowledge | Understanding Science Content | Student displays the knowledge and understanding of a scientific concept, idea, skill, or practice | *"I didn't know anything about preprints before this class. It was also a bit surprising to hear that you can cite these papers. This could be especially helpful because, if the publishing process is slower than ever, it would take longer for active projects to hear about work being done in conjunction. But, through preprints, they have the opportunity to keep the science even more up to date."* |
| | | Engaging in science discourse | Student shares how they have initiated conversations about science in and/or out of class | *"My grandmother was just as shocked as me the we have to pay to have our reviews published and it was really lovely to be able to share my education with my family and have them listen to my thoughts."* |
| | Practice | Application of science skills/knowledg e | Student shares their ability to practice the skills and knowledge that they have learned | *"After thoroughly reviewing the standard requirements for a publication, as well as the scientific theory supporting the article, I felt surprisingly well-equipped to offer constructive feedback on the assigned pre-print."* |
| | Value | Understanding the use and need of science | Student discusses importance of science knowledge and/or practice within the science community or to broader society | *"... it [peer review] gives different perspectives and each person is going to find different things. Because the reviewers will have slightly different specialties in terms of their own research, they will be able to pick out different things that may not make sense to someone outside a niche field or give feedback on a technique that they have extensive knowledge about that is done well/could be optimized for better results."* |
| Identity | Personal Identity | Perceived advantages | Student shares how one or more of their personal identities have positively | *"...when I consider my class identity and familial background in relation to the research process as a whole (I'm including peer* |

| | | | | |
|---|---|---|---|---|
| | | | impacted their ability to pursue science | *review in this process), I know that my understanding and appreciation for this process comes from being supported and taught about the significance of information, as well as having the social capital to see myself involved in that process. "* |
| | | Perceived disadvantages | Student shares how one or more of their personal identities have negatively impacted their ability to pursue science | *"I am a chicane disbaled queer trans person who is in stem. People like me just do not exist in stem and if they do its only one person and often time they do not share all of my identities. This experience will always impact me as I am always the only person who will look like me in the room and I have just come to terms with it at this point which is something that is sad but what can I do."* |
| | Professional Identity | Pursuing science career/internships | Student discusses their current or future plans as a professional scientist | *"...it [participating in peer review] gives me something to add to my CV as well as show my participating in the science world that is beyond my classes"* |
| | | Interacting with science professionals | Student shares how interacting with professional scientists solidifies their perception of themselves as a competent scientist | *"... communicating with authors about manuscripts would be extremely beneficial in really envisioning myself as a scientist."* |
| Belonging | Presence | | Student explicitly states that they feel a sense of belonging within the science community OR mentions feeling welcomed/validated as a scientist | *"Working on the peer review made me feel like I was making a valuable contribution to the scientific community and for me to do so I felt like I am a knowledgeable participant."* |
| | Absence | | Student explicitly states that they do NOT feel a sense of belonging within the science community | *"Doing review #1 did not strengthen the feeling of belongingness to the science community because I am not sure of so many things in my review. I constantly use" might be" and "possible" in sentences to leave space for correctness."* |

| | | | | |
|---|---|---|---|---|
| | Facultative | | Student talks about spaces where they do and don't feel belonging | *"In the past, I have done journal clubs with graduate students, which have been much more difficult for me to feel confident in sharing my thoughts. Today's class really helped me gain more confidence about my ability to review primary research independently."* |
